# Chromosome assembly for the Black bean aphid *Aphis fabae*

**DOI:** 10.64898/2026.08.05.743085

**Authors:** Mark Whitehead, Claudia Wierzbicki, Margaret Hughes, Alistair Darby

## Abstract

The black bean aphid, *Aphis fabae* is a crop pest and vector of insect-transmitted pathogens, comprising closely related sub-species with overlapping host ranges. In other *Aphis* species, over-expression of specific detoxification genes has been linked to insecticide tolerance. We present two chromosome-scale assemblies for a clonal *A. fabae* line, representing two phased haplotypes, generated using HiFi and Hi-C sequencing technologies. A comprehensive genome annotation, built with PacBio Iso-Seq data, was used to investigate genes underlying insecticide tolerance. Both genomes are comprised of four chromosomal blocks (haplotype 1: 427 Mb; haplotype 2: 396 Mb) with high BUSCO completeness (98.7%). Comparative genomics revealed an expansion of UDP-glycosyltransferases, whose expression is linked to insecticide detoxification in other *Aphis* species. These high-quality references provide a foundation for studying *A. fabae* sub-species and a genomic resource for investigating insecticide tolerance across the *Aphis* genus.

**Author summary:** Here we have provided a comprehensive assembly and annotation for further study into the Black bean aphid, *Aphis fabae*, using up to date long-range sequencing technologies. The final assemblies for both haplotypes are chromosome length and consist of 4 main chromosome blocks, consistent with the literature. The *A. fabae* genome was found to contain an increase in copy number of UDP-glycosyltransferases, which have previously been linked to insecticide resistance. The work here will be a resource to those studying insecticide tolerance in crop pests, as well as the differences between *A. fabae* sub-species.

## Introduction

The black bean aphid, *Aphis fabae,* is a polyphagous aphid pest found on ornamentals and economically important crops such as broad bean and sugar beet[1]. *A. fabae* is also considered a group of very similar aphid sub-species that have overlapping primary and secondary hosts[2][3]. Previous work has gone into defining the different sub-species of *A. fabae*, along with changes to taxonomic changes their classification, such as *Aphis evonymi*, now it’s own species while still sharing *A. fabae* characteristics[4][5][6][7].

Aphids are such a successful pest due to their ability to proliferate rapidly during Spring and Summer. After overwintering as eggs on a primary host plant, aphids undergo cyclical parthenogenesis (CP), generating multiple generations of clonal offspring in a short space of time[8]. Insecticide resistance may also play a role in aphid survivability. Mechanisms of resistance and their impacts are summarised in Bass & Nauen (2023)[9], with resistance to acetylcholinesterase (AChE) inhibitors and higher levels of esterase activity often seen as disruptive to Carbamate and Organophosphates class insecticides[10][11][12]. Species belonging to the *Aphis* genus, such as *Aphis craccivora* and *Aphis gossypii*, are observed having increased insecticide tolerance[13][14][15], the latter of which is linked to the expression of uridine-diphosphate (UDP)-glycosyltransferases (UGTs). UGTs facilitate the excretion of toxic compounds by transforming endogenous molecules into a more hydrophilic state, allowing easier elimination from the body[16]. UGT resistance is also linked to medically important disease vectors, so their study may have implications further than agricultural settings[17][18]

Since the first aphid genome for *Acyrthosiphon pisum* was published in 2010[19], there has been a large expansion in the number of aphid genomes. As of 2025, the NBCI genomes portal has 74 Aphididae genomes, of which 24 are at chromosome level, comprising of 41 unique species. Chromosome level assembly is generally only achievable through the advent of both long accurate PacBio high fidelity (HiFi) reads [20], and the ability to identify sequence proximity spanning megabases at time while improving haplotype phasing with Hi-C chromosome capture[21][22]. Alongside developments in genome assembly, we also see improvement in transcriptome generation using PacBio Iso-seq technology and Oxford Nanopore cDNA sequencing[23][24]. Not only do these methods permit full length transcript sequencing, they also help in novel isoform discovery, both of which are often difficult for short read methods, and can result in a greater understanding of a range of biological mechanisms and pathways[25][26][27]. These high molecular weight sequencing technologies have been used to facilitate high-quality genomes for aphids such as *Sitohbian avenae*[28], *A. gossypii*[29], *Rhopalosiphum maidis*[30], as well improved assemblies for *A. pisum*[31] and *M. persicae*[10].

Here, we use a combination of current state of the art technologies in high molecular weight sequencing to generate two haplotype assemblies for *A. fabae,* both of which are at chromosome level. We utilise long read Iso-seq data for genome prediction and annotation, which we then interrogate for the presence of detoxification genes commonly linked to tolerance to insecticidal compounds while demonstrating the increased count of UGT’s in *A. fabae*. This work will be a resource for further Aphididiae comparative genomics, as well as provide a basis for further work into insecticide resistance and the *A. fabae* species complex.

## Methods

### Sampling

Aphids were sampled a private garden in Meols, Wirral. A single healthy aphid was selected to start a clonal line consisting of aphids which would therefore be genetically identical. Aphids were maintained on whole bean plants in a climate-controlled incubator at 16 °C, 16L:8D (light dark cycle) and 60% relative humidity.

### DNA/RNA extraction

DNA was extracted from 100 aphids using a standard phenol-chloroform extraction using phase-lock tubes to transfer aqueous solution, which helps reduce the effect of sheering forces on high molecular weight DNA which are normally introduced using pipette transfer[32]. For contig orientation into chromosomes, aphids were flash frozen in liquid nitrogen and sent to Phase Genomics (Seattle, USA) for Hi-C sequencing. We produced five total RNA extractions from samples consisting of either nymphs, five day old apterous juveniles, five day old alate juveniles (based on the presence of developing wing buds), winged adults, and finally a mix sample consisting of the remaining culture. Using a range of morphs and life-stages helps capture differentially expressing genes across the aphids life cycle. RNA extraction was performed using Zymo-quick RNA kit (California, USA) following the manufacturers instructions.

### Whole genome sequencing

5ug of genomic DNA was sheared to a suitable size for HiFi using the Megaruptor 3 system. The size was checked with the fragment analyser. This sample was the direct input for removal of single stranded overhangs at 37°C for 15 minutes, DNA damage repair step at 37°C 30 minutes and an end repair step at 20°C for 10 minutes and 65°C for 30 minutes. Overhang adapters were added in a 20°C step for 1hour. After this the sample was cleaned with 1:1 ampure beads. The sample was then subjected to an enzyme clean step at 37°C for 30 minutes followed by a further ampure clean step as before. The sample was size selected using a 0.75% cassette and s1 marker on the SAGE blue pippin system using the range 8-50kb. The recovered sample was cleaned with 1:1 ampure beads and checked on the fragment analyser. The sample was bound to Sequel Polymerase 2.2 and sequenced on the PacBio Sequel IIe using Adaptive loading and 30h movie time.

### RNA Iso-seq sequencing

Iso-Seq libraries were prepared for sequencing using the SMRTbell® prep kit 3.0. Samples were barcoded during cDNA amplification and amplified for a total of 15 cycles. The barcoded libraries were equimolar pooled. The SMRTbell library was prepared for sequencing using Sequel II binding kit 3.1 and sequenced on a single SMRT cell on the PacBio Sequel IIe with 24-hour movie time.

### Genome assembly

Genome characteristics were assessed using Genomescope (v2)[33]. Initial contig assemblies were generated using hifiasm (v0.16.1-r375)[34] and was provided with both HiFi and Hi-C data for improved haplotype resolution. Two chromosome phased genome assemblies are yielded from hifiasm, denoted as haplotype 1 and haplotype 2. Both genome copies were individually scaffolded into linkage groups using YaHS (v1.2a.2)[35]. Blobtools (v1.1)[36] was used to identify symbiont contigs present in both assemblies, which were subsequently removed. Assembly completeness was assessed using BUSCO (v5.2.2)[37] and the arthropoda_odb10 database. The putative X chromosome (scaffold_3 and scaffold_5) for assembly the haplotype 1 assembly (hap1) was not scaffolded into a single chromosome. This was corrected using RagTag (v2.0.1)[38] and scaffold_1 from the hap2 assembly, where the X chromosome consisted of a single scaffold.

### Accessory genomes

The original aphid HiFi read set was assembled using Hifiasm and default settings, but with --primary enabled. Bacterial genomes and any associated plasmids were identified manually using blobtools and megablast outputs. Bacterial genomes were then annotated using bakta (v1.9.3)[39]. HiFi reads were also provided to MitoHiFi (v3.0.0)[40] for mitochondrial genome assembly and annotation. We identified genomes for the obligate aphid symbiont *Buchnera aphidicola,* a *Rickettsia* species identified as *R. viridis* based on online NCBI Blast results of the 16S sequence, as well as a *Pantoae* species. No further analysis was carried out on these genomes, but have been made available in the bioproject resource (supplementary table 1).

### Genome annotation

Each genome copy was softmasked using Repeatmasker (v4.0.7)[41] and custom-built repeat model libraries using RepeatModeler (v.1.0.11)[42]. Iso-Seq reads were demultiplexed using PacBio SMRT-link (v13) which provides a “hq_transcripts.fa” file of high-quality transcripts. High-quality transcripts were mapped to the genome using minimap2[54]. For each genome copy assembly, transcript assemblies were initially generated using PASA (v2.5.2)[43] with hq_transcripts.fa as input. PASA transcript assemblies were then provided to Augustus (v3.5.0)[44] for model training and gene prediction. The outputs from PASA, Augustus, mapped transcript alignments and mapped protein sets (Swiss-prot proteins and orthoDb Arthropoda v10 proteins) using miniprot (v0.11-r234)[45] were provided for consensus gene modelling with EvidenceModeler (v2.0.0)[46]. Transposable elements were identified and removed via diamond blastp (v2.0.15)[47] against the TransposonPSI database (https://transposonpsi.sourceforge.net/). Gene models were updated with untranslated regions (UTRs) and alternate isoforms using PASA update. Completeness was assessed using BUSCO against the *A. fabae* proteome with the Arthropoda odb10 database.

### Comparative genomics

Protein fasta sequences for other Aphid species were downloaded for comparative study. These aphid species and their download locations are described in supplementary table 2. For phylogenetic tree inference, single copy orthologues were identified across all these species using orthofinder (v2.5.4)[48]. Proteins within each orthogroup were aligned using mafft (v7.487)[49], with alignments then trimmed using gblocks (v0.91b)[50]. Alignments were concatenated and used as input for model selection using modeltest-ng (v0.1.7)[51]. Finally, the resulting selected model and alignment was provided to raxml-ng (v0.6.0)[52] (10 initial starting tree and 1000 bootstrap replicates) for tree inference. The final tree was visualised using the Interactive Tree of Life online tool (https://itol.embl.de/).

Synteny of the four *A. fabae* chromosomes was assessed against the chromosome scale assembly for *A. gossypii*[29]. We identified arthropod single-copy orthologues in both genomes using BUSCO. Collinearity of BUSCO genes between both genomes were then identified using MCScanX (v1.0.0)[53] with default settings and subsequently visualised with the Synvisio online tool (https://synvisio.github.io/).

## Results and discussion

### Assembly and Annotation

Using a combination of PacBio and Hi-C chromosome linkage information is key to generating high-quality chromosome assemblies. Not only that, but the use of Hi-C will also allow us to phase each chromosome, reducing assembly mosaicism (sequences consisting of haplotype blocks from different chromosome sets). A total of 13.1 Gb of HiFi data equating to approximately 33x coverage was generated. We were provided with 78.9 Gbp of Hi-C reads from phase genomics.

Assembly statistics for both genome copies are available in table 1. The primary assembly is both highly contiguous (n50=97.5 Mb) and complete based on BUSCO scores (98.5% complete with Arthropod BUSCO genes). We yielded four chromosomes (figure 1), with 91.5% of the assembly sequence present within these four chromosome sequences. The final assembly is also in-keeping with the predicted genome length from genomescope analysis (371.0 Mb). The alternate reference is also very complete (98.6% complete with Arthropod BUSCO genes), but also comprised of more small debris contigs, as well as a larger overall assembly size of 426.8 Mb. When we compare the primary assembly to other available Aphididae, we observe the *A. fabae* assembly is consistent with other chromosome level assemblies regarding length and n50, while also being as complete based on BUSCO scores (supplementary figure 1).

**Figure 1.**
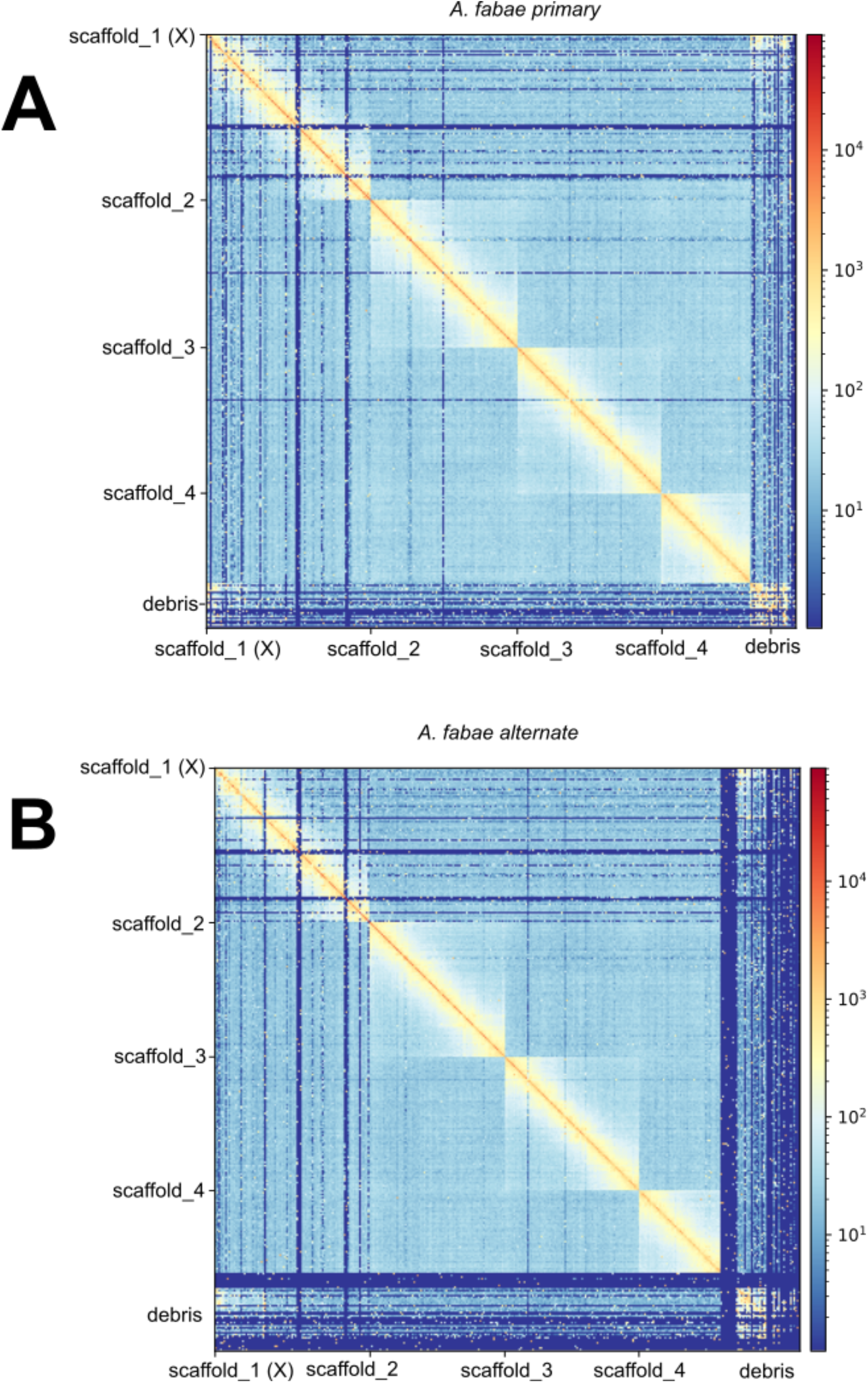
Hi-C contact maps primary and alternate assemblies of *A. fabae*. Both assemblies consist of four chromosomes as suggested by the literature. The alternate haplotype assembly contains more debris sequence, while also requiring manual curation of the putative X chromosome.

**Table 1.** Genome assembly statistics.

|  | ***A. fabae* primary** | ***A. fabae* alternate** |
| --- | --- | --- |
| **genome length (bp)** | 395,917,334 | 426,774,156 |
| **max length (bp)** | 109,341,639 | 111,241,487 |
| **min length (bp)** | 2,000 | 11,000 |
| **num. scaffolds** | 97 | 193 |
| **Num. Ns (bp)** | 4,600 | 3,300 |
| **GC%** | 27.73% | 28.11% |
| **n50 (bp)** | 97,553,529 | 96,326,046 |
| **BUSCO** | C:98.7%[S:95.8%,D:2.9%],F:0.5%,M:0.8%,n:1013 | C:98.6%[S:95.7%,D:2.9%],F:0.5%,M:0.9%,n:1013 |

Initial repeat-masking of the assembly prior to annotation identifies 29.5% of the genome belonging to repetitive sequence (supplementary table 3). While 20.81% belongs to unclassified repeats, the next largest group were DNA transposons, which account for 5.44% of all identified repeats. A soft-masked version of the assembly was used gene prediction. We used a custom approach to genome annotation, as opposed to existing pipelines which are based on illumina short-read RNA-seq evidence. The benefit of Iso-seq here will allow discovery of transcripts unidentifiable with short reads alone, which may be unable to span multiple exons across a transcript. We predicted 18,873 protein coding genes, which in-turn encode for 25,814 mRNA transcripts. The proteome is very complete when looking at BUSCO scores (C:96.2%[S:63.9%,D:32.3%],F:1.2%,M:2.6%,n:1013, arthropoda_odb10). Additionally, we further add putative functional annotations with interproscan, eggnog and SignalP, which annotate 15,027, 21,375 and 1,888 proteins respectively.

### Comparative genomics

*A.* fabae and *A. gossypii* chromosomes share high levels of synteny (figure 2), with 96.38% of all single-copy arthropod BUSCO genes shared between both species. The *A. fabae* X chromosome (scaffold_1) was identified through homology to the X chromosome of *A. gossypii* (NC_065533.1).

**Figure 2.**
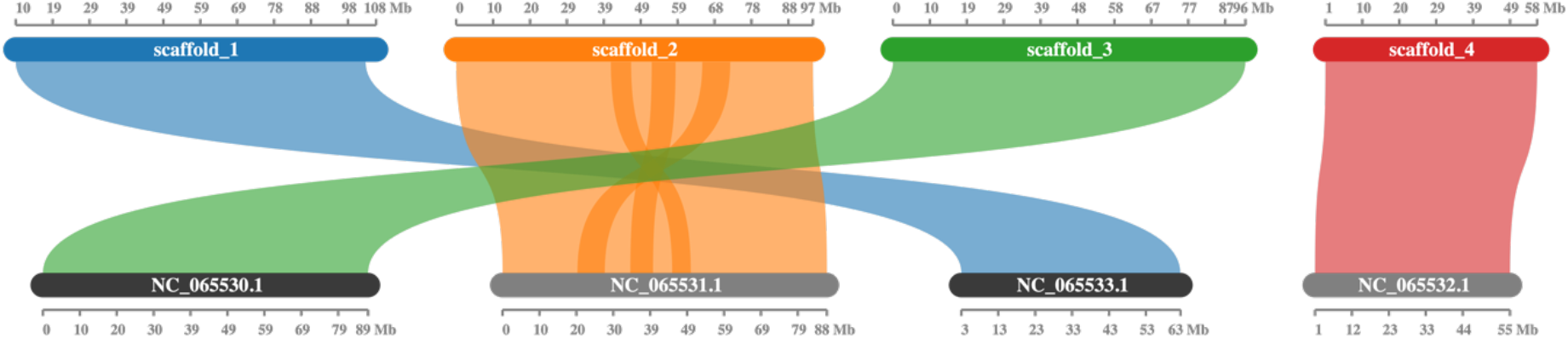
Chromosome synteny between *A. fabae* (top) and *A. gossypii* (bottom). Sequences from each organism are ordered via length. Homology of NC_065533.1 suggest the X chromosome to be scaffold 1. We see a high level of shared synteny across the chromosomes, which is to be expected based on their phylogenetic relatedness.

We identified a large break in the X chromosome of haplotype assembly 1. We did not see this same break in primary assembly, therefore we provided the unbroken primary assembly chromosome X to ragtag to scaffold together the two contigs that comprise the X in the alternative assembly. In both haplotypes, there are large gaps on the X where there are little-to-no Hi-C contacts present. These could be repetitive regions or inactive and inaccessible heterochromatin, which results in regions of little access for Hi-C contacts to be generated.

Based on phylogenetic placement (figure 3A), we identified shared orthogroups as identified with Orthofinder between *A. fabae* and three other related aphid genomes; *A. gossypii*, *R. maidis* and *A. glycines* (figure 3B). While all four shared 7,880 orthogroups, both *A. fabae* and *A. glycines* had a high number of gene families unique to them (314 and 322 respectively).

**Figure 3.**
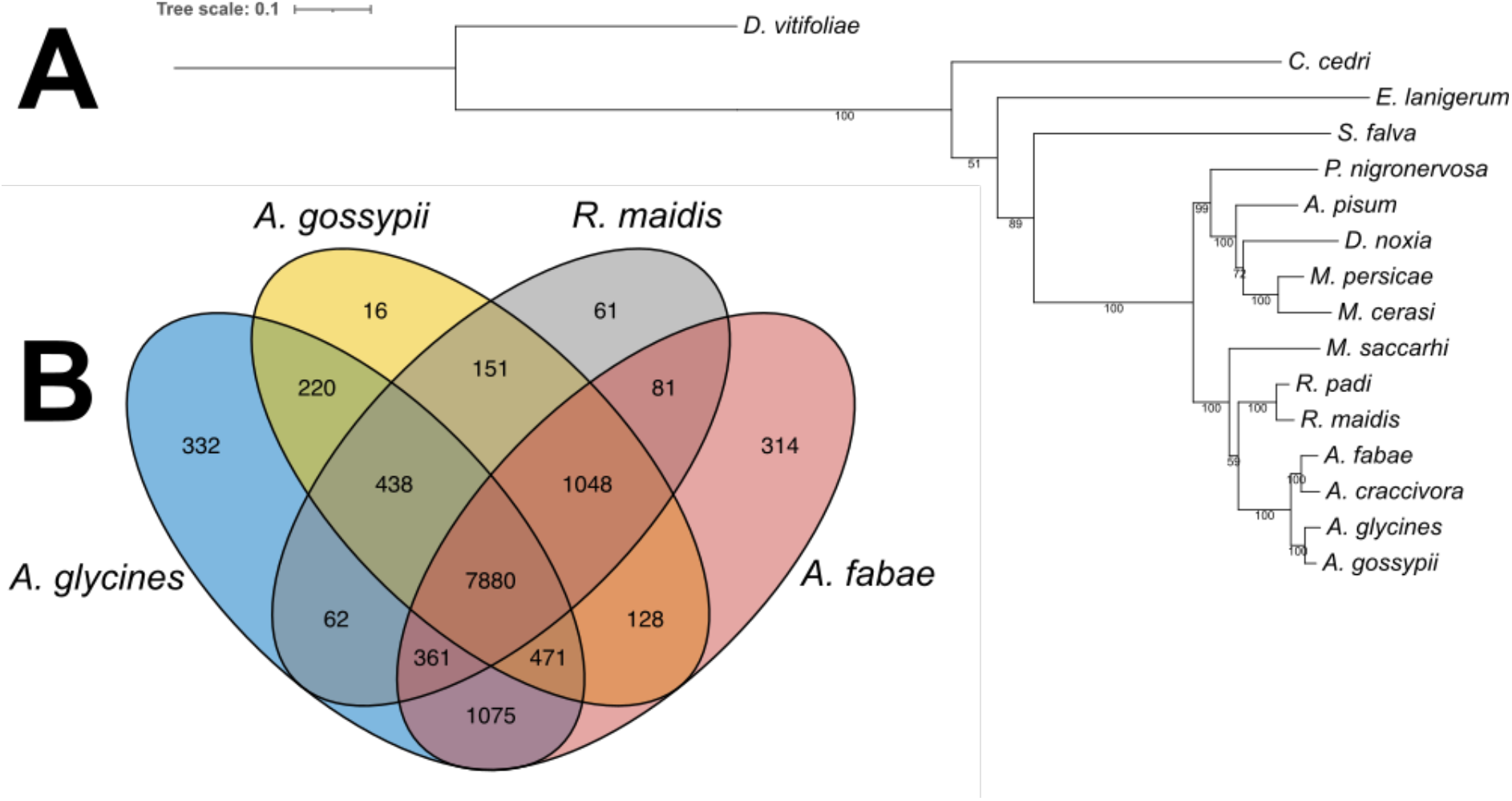
Aphididae comparative genomics. A) Phylogenetic analysis of aphid species based on 82 shared single copy orthologues across all species using *D. vitifoliae* as an outgroup. The tree was created using a maximum-likelihood method with 1000 bootstraps. Node numbers indicate support values. Scale bar indicates amino acid substitutions per site. JTT+I+G4 was the amino acid substitution; B) Four-way venn diagram demonstrating shared gene orthologue groups between *A. fabae*, *A. gossypii*, *R. maidis* and *A. glycines*.

Genes associated with detoxification functions are commonly linked to insecticide/pesticide tolerances. We assessed the gene copy frequencies for cytochrome p450 (cP450), glutathione-S tranferases (GSTs), carboxyl esterases (Cbxyl-ests), UDPGs, and ABC transporters (ABCts). cP450, GST and ABCt counts were similar across all four species, while Cbxyl-est counts were slightly elevated in *A. fabae*. The increase in 65 UDPG genes was more apparent, by having 14 more copies than the next highest count (51 in *A. gossypii*). UDPG’s have previously been implicated in insecticide tolerance in *A. gossypii*. The increase of UDPGs in *A. fabae* suggests again that they play a major role in detoxification of foreign bodies, such as pesticides.

**Table 2.** Detoxification gene frequencies across *A. fabae*, *A. gossypii*, *R. maidis* and *A. glycines*.

| function | *A. fabae* | *A. glycines* | *A. gossypii* | *R. maidis* |
| --- | --- | --- | --- | --- |
| **cP450** | 56 | 58 | 56 | 47 |
| **GSTs** | 11 | 11 | 9 | 9 |
| **Cbxyl-est** | 29 | 25 | 21 | 18 |
| **UDPGs** | **65** | 49 | 51 | 41 |
| **ABCts** | 69 | 70 | 71 | 67 |

## Conclusions

Here we report two high quality assemblies for *A. fabae* representing primary and alternate haplotype assemblies for all chromosome copies. With *A. fabae* being a vector of plant and crop disease, the genomes presented here will be useful for further understanding of possible insecticide tolerances, especially regarding the role of UDPGs. There is also scope for further work into the population structure and sub-types of *A. fabae*. While most genomes available up until now are haploid assemblies, consisting of a mosaic of all chromosome copies, here we ameliorate mosaicsim using newer HiFi and Hi-C technologies. We also demonstrate the use of Iso-seq data for gene prediction and their ability to yield more gene isoforms.

## Supplementary

**Supplementary figure 1.**
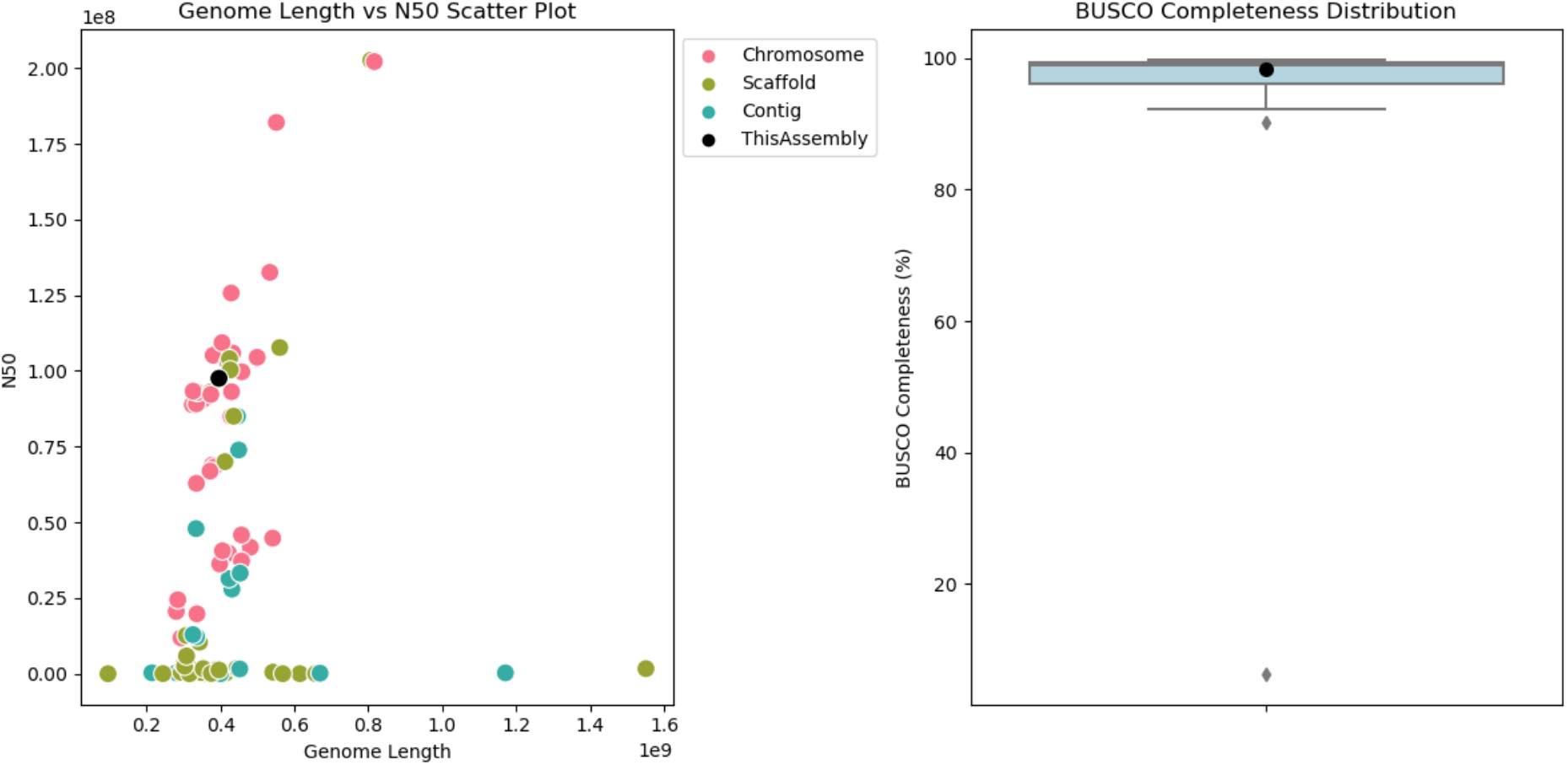
Genome completeness in *A. fabae* primary assembly compared to other Aphididae species. Genome statistics were pulled using the Genome on a tree (GOAT) tool. BUSCO scores are based Eukaryotic orthologue databases.

## Data availability

All reads and assemblies are submitted to ENA. Accessions for all data can be found in supplementary table 1.

## Acknowledgments

Sequencing was performed by the Centre for Genomic Research (CGR) at University of Liverpool. We would also like to thank Lukasz Lukomski for finding the *Aphis fabae* sample from which we derived the clonal line used in this study.

## Supplementary tables

See supplementary_tables_27Jan.xlsx

- **Supplementary table 1.** Bioproject associated accessions.
- **Supplementary table 2.** Download locations for protein fastas used in phylogeny construction.
- **Supplementary table 3.** Repeatmasker summary table for A. fabae (primary assembly).

